# Antigen specificity shapes antibody functions in tuberculosis

**DOI:** 10.1101/2024.06.03.597169

**Authors:** Joshua R. Miles, Pei Lu, Shuangyi Bai, Genesis P. Aguillón-Durán, Javier E. Rodríguez-Herrera, Bronwyn M. Gunn, Blanca I. Restrepo, Lenette L. Lu

**Affiliations:** UT Southwestern Medical Center, Department of Immunology; UT Southwestern Medical Center, Division of Infectious Diseases and Geographic Medicine, Department of Internal Medicine; Paul G. Allen School of Global Health, College of Veterinary Medicine, Washington State University; Department of Epidemiology, School of Public Health, University of Texas Health Science Center at Houston, Brownsville campus, Brownsville, TX, USA; Departamento Estatal de Micobacteriosis, Secretaría de Salud de Tamaulipas, Reynosa 88630, Matamoros 87370, Tamaulipas, México; School of Medicine, South Texas Diabetes and Obesity Institute, University of Texas Rio Grande Valley, Edinburg, TX, USA; I.CARE and Population Health, Texas Biomedical Research Institute, San Antonio, TX, USA; Parkland Health

## Abstract

Tuberculosis (TB) is the number one infectious disease cause of death worldwide due to an incomplete understanding of immunity. Emerging data highlight antibody functions mediated by the Fc domain as immune correlates. However, the mechanisms by which antibody functions impact the causative agent *Mycobacterium tuberculosis (Mtb)* are unclear. Here, we examine how antigen specificity determined by the Fab domain shapes Fc effector functions against *Mtb.* Using the critical structural and secreted virulence proteins *Mtb* cell wall and ESAT-6 & CFP-10, we observe that antigen specificity alters subclass, antibody post-translational glycosylation, and Fc effector functions in TB patients. Moreover, *Mtb* cell wall IgG3 enhances disease through opsonophagocytosis of extracellular *Mtb*. In contrast, polyclonal and a human monoclonal IgG1 we generated targeting ESAT-6 & CFP-10 inhibit intracellular *Mtb*. These data show that antibodies have multiple roles in TB and antigen specificity is a critical determinant of the protective and pathogenic capacity.

## Introduction

While antibodies are leveraged for vaccines, diagnostics, and therapeutics in many infectious diseases, their role in tuberculosis (TB) is unclear (*1–9*). Cellular immunity is thought to be the cornerstone of protection (*3, 10*). However, enhanced T cell responses through immune checkpoint inhibitors paradoxically worsen disease (*11, 12*). Moreover, the first large phase IIB clinical trial of the TB vaccine MVA85A designed to boost Th1 and Th17 responses showed no protection (*13*). Subanalyses showed that CD4 T cell responses associated with increased whereas Ag85A IgG linked to decreased risk of disease. Thus, antibodies have the potential to modulate TB and understanding the mechanisms by which antibody functions impact *Mtb* pathogenesis would enable us to leverage this potential more effectively.

Antibodies use the Fab domain to recognize antigen and Fc domain to elicit immune cell effector functions (*2, 5*). Fc domain diversity in subclass and post-translational glycosylation regulate binding to activating and inhibitory Fc receptors (FcRs) that initiate downstream signaling and immune cell effector functions (*14–17*). In mice, loss of activating FcR signaling decreases survival after *Mtb* challenge (*18*). Conversely, loss of inhibitory FcR signaling leads to decreased bacterial burden. In humans, the activating FcγRIIIa is associated with protection in latent TB where bacterial burden is undetectable. This contrasts disease in active TB where *Mtb* is identifiable, transmissible, and lethal if untreated (*19*). We have shown that IgG Fc properties and effector functions diverge in latent and active TB (*20, 21*). Moreover, treatment with IgG from latent compared to active TB leads to decreased *Mtb* burden and increased antimicrobial activities in an *in vitro* macrophage model of *Mtb* infection (*20*). These findings suggest that polyclonal antibodies in TB patients differ in the capacities to modulate *Mtb* infection but whether they are protective or disease enhancing remains to be defined.

Antigen specificity as determined by the Fab domain impacts Fc effector functions. Changes in the Fab domain alters the ability of the antibody Fc domain to bind to FcRs and initiate effector functions. For influenza and Cryptococcus, this change is sufficient to alter mAb mediated protection (*22–24*). For TB, mAbs that target different *Mtb* surface and cell wall antigens (LAM, PstS1, and heparin-binding hemagglutinin are examples) utilize different FcRs to impact *Mtb* (*25–28*) but what occurs in latent and active TB is not known. To understand how the target of the antibody modulates the ability of the Fc domain to alter *Mtb* pathogenesis, here we use unique *Mtb* antigens to dissect polyclonal antibody Fc effector functions in TB patients.

## Results

### Antigen specificity alters the capacity of antibodies to differentiate latent and active TB

We used patient samples from well characterized latent TB infection and active TB disease because these states in the clinical TB spectrum link to protection and disease (Table) (*29*). Groups were matched in age, sex, and Bacillus Calmette-Guérin (BCG) vaccination. Individuals with co-morbid conditions such as HIV and type II diabetes were excluded. Endemic TB negative controls were used to account for exposure to environmental non-tuberculous mycobacteria and other geographical variations that impact immunity (*30–34*).

To systematically approach the >4000 *Mtb* protein antigens (*35*), we balanced breadth and specificity with individual and fractions of mixed antigen preparations from bacteria in culture. Since the dominant repertoire of antigens recognized by antibodies in TB is unclear (*36*), many studies use purified protein derivative (PPD) which is mixture of proteins from bacterial culture (*6, 37, 38*). In addition to PPD, we used protein fractions enriched in components of *Mtb* cell wall, cytosol, and secretions into the culture (culture filtrate) (*39–41*) on the premise that localization of the antigen could impact the sensing, processing, and development of antibodies (*26, 27, 42–45*). For specific proteins, we chose the well-studied secreted bacterial virulence factor complexes Ag85A & Ag85B (*46*) and ESAT-6 & CFP-10 (*47*). As a non-*Mtb* control from a common pulmonary pathogen with high seroprevalence in adults, we used respiratory syncytial virus (RSV) (*48*). Together these antigens enabled the evaluation of humoral immunity targeting bacterial structural, metabolic, and immune modulatory functions important for pathogenesis.

We first examined the impact of antigen specificity on how IgG titers relate to infection and disease. Only *Mtb* cell wall IgG could distinguish between latent and active TB (Figure 1A), primarily by IgG1 and IgG2 (Figure 1B). In contrast, IgG reactive to PPD, *Mtb* cytosolic proteins, culture filtrate, and Ag85A & Ag85B and ESAT-6 & CFP-10 could not distinguish between clinical TB states (Figure 1A and 1C). ESAT-6 & CFP-10 IgG was even worse than control RSV IgG (Figure 1A, 1C and Supplemental Figure 1). These data are consistent with the poor ability of *Mtb* reactive IgG in serodiagnostics to identify clinical TB (*49, 50*). We focused on *Mtb* cell wall and ESAT-6 & CFP-10 to understand the antibody Fc effector functions linked to IgG reactive to antigens with the most and least capacities to differentiate latent and active TB.

**Figure 1.**
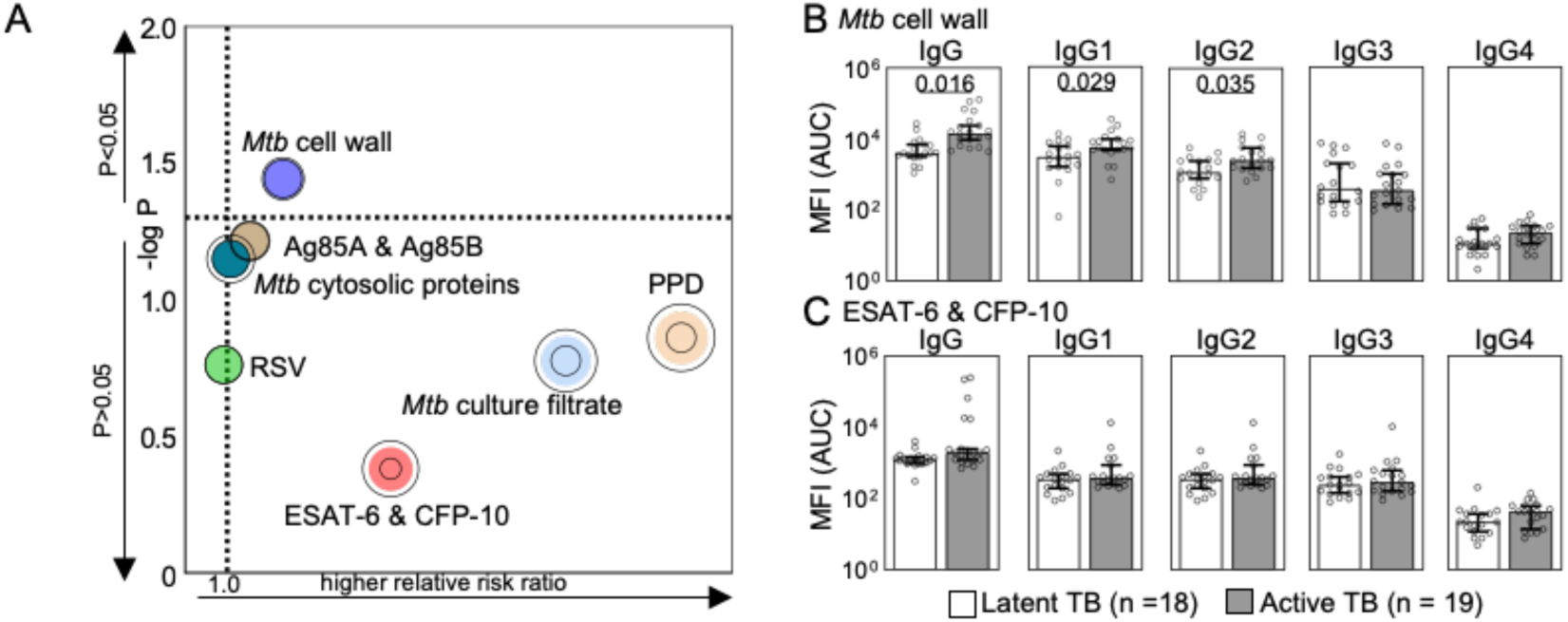
Antigen specificity changes how IgG associates with latent and active TB. **(A)** The bubble plot shows the capacity of each antigen specific IgG to predict active TB disease and latent TB infection by relative risk ratios (RRR) and 95% CI (rings), with higher likelihood of active TB >1 and latent TB <1. The median and 95% CI of IgG reactive to **(B)** *Mtb* cell wall and **(C)** ESAT-6 & CFP-10 are shown with *P*-values adjusted for sex and age.

### Antigen specificity drives differential antibody Fc domain glycosylation in latent and active TB

Antibodies function through diversity of the Fab domain via its antigenic repertoire and the Fc domain from subclass distribution and post-translational N-glycosylation (*2, 14–17*). On a single conserved residue on the IgG Fc domain (N297) is a core biantennary complex of mannose and N-acetylglucosamine (GlcNAc) (Supplemental Figure 2A). Addition and subtraction of galactose, sialic acid, bisecting GlcNAc, and fucose to the core structure generates heterogenous glycoforms. For every individual cohort sample, we measured the glycoforms on IgG reactive to ESAT-6 & CFP-10, *Mtb* cell wall, control RSV, and total bulk IgG Fc domains (Figure 2A, Supplemental Figure 2B) and summarized them (Figure 2B, Supplemental Figure 2C and 2D). We observed that antigen specific differed from total bulk IgG glycans with higher sialic acid, galactose, and bisecting GlcNAc (Figure 2B and Supplemental Figure 2D). Between antigen specific IgG, *Mtb* antigens distinguished themselves from RSV through even higher sialic acid, galactose, and bisecting GlcNAc and lower fucose (Figure 2B and Supplemental Figure 2D). Between *Mtb* antigens, *Mtb* cell wall compared to ESAT-6 & CFP-10 IgG had lower sialic acid and galactose and higher fucose, with the major changes occurring in fucose without sialic acid. These data show that each antigen specificity has a unique glycosylation pattern, consistent with B cell intrinsic rather than extrinsic mechanisms of IgG glycosylation (*51, 52*).

**Figure 2.**
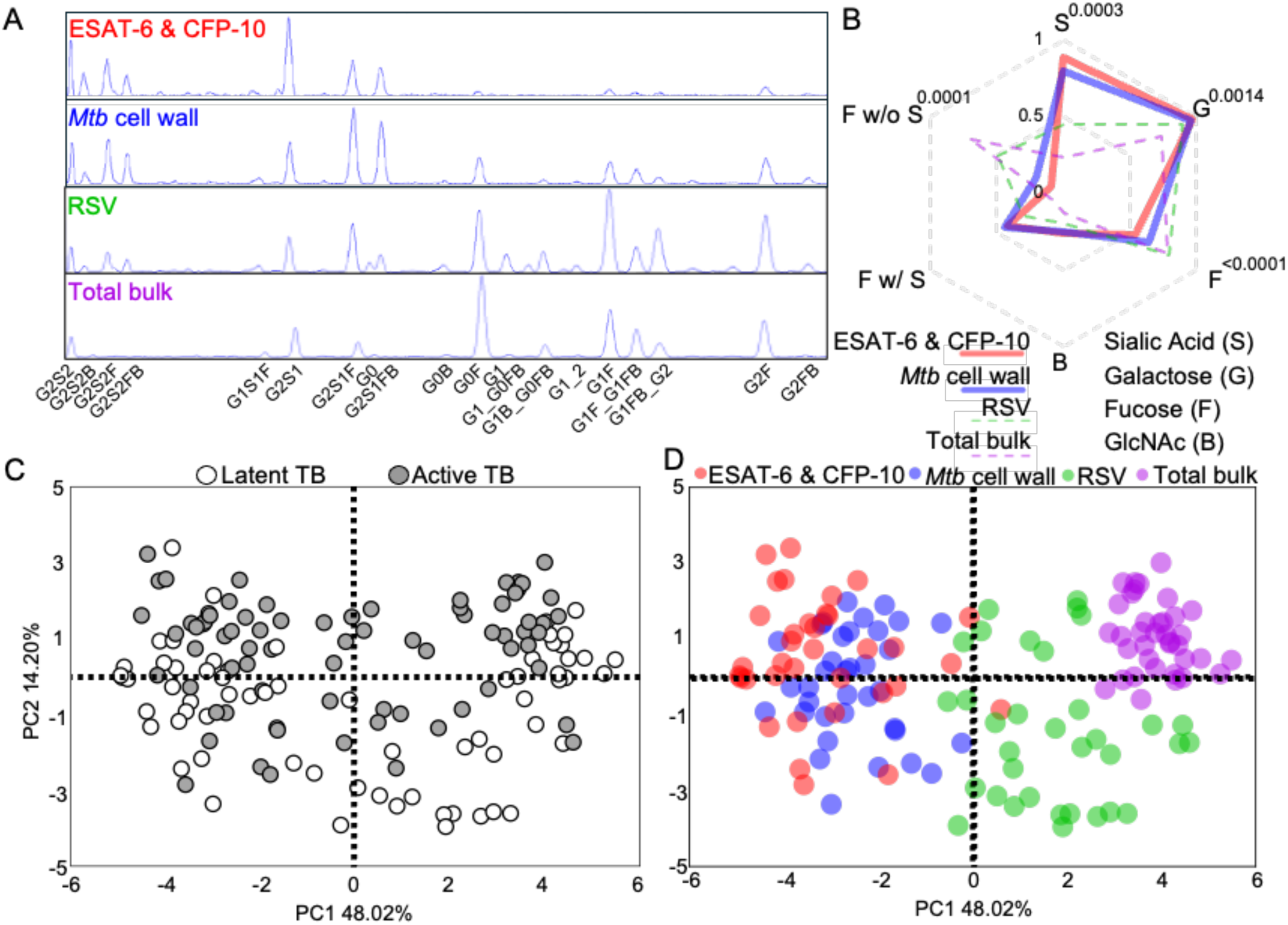
Antigen specificity separates antibody Fc domain glycosylation in latent and active TB. **(A)** Representative chromatograms show the patterns of individual glycoform isolated from the Fc domain of antigen specific and bulk total IgG in an individual TB patient sample. **(B)** The radar plot shows the relative abundance of total glycans from antigen specific and bulk total IgG with lines depicting the median of all TB patient samples. Comparisons between ESAT-6 & CFP-10 and *Mtb* cell wall IgG glycans were made by Wilcoxon matched-pairs signed-ranks tests and significance after adjustment for multiple comparisons is shown. **(C – D)** Principal components analysis was performed. Each dot summarizes the individual glycoforms for ESAT-6 & CFP-10, *Mtb* cell, RSV, or total bulk IgG for each individual TB patient. Score plots show clustering of glycosylation patterns with respect to **(C)** latent and active TB and **(D)** antigen specificity.

We previously reported that IgG glycosylation diverges between latent and active TB (*20, 21, 53–55*). While differences between these two clinical states were observed here (Figure 2C), antigen specificity had a greater impact (Figure 2D and Supplemental Figure 2F) in principal components analysis and hierarchical clustering. Thus, antibody glycosylation is influenced by antigen specificity more than clinical TB state.

### Antibody Fc effector functions diverge more by antigen specificity than latent and active TB

We hypothesized that differences in subclass distribution and Fc N-glycosylation between *Mtb* cell wall and ESAT-6 & CFP-10 IgG indicated divergence in Fc receptor (FcR) binding and immune cell effector functions. We tested this hypothesis by evaluating the ability of antigen specific IgG in each individual TB patient to engage FcRs described to impact TB: FcγRI (*25, 56, 57*), FcγRIIa (*25, 26, 43*), FcγRIIb (*18, 25, 26, 43*), FcγRIIIa (*19, 20, 26*), and FcRn (*27, 58*) (Supplemental Figure 3A). To incorporate adaptor-mediated signaling and feedback, we used the cell-based assays antibody dependent natural killer cell activation (ADNKA) that induces cellular cytotoxicity and antibody dependent cellular phagocytosis (ADCP) (Supplemental Figure 3B and C), both of which are differentially engaged in latent and active TB (*19, 20, 25, 53*). We found that Fc effector functions were defined by antigen specificity more than TB status (Figure 3A and 3B). ADNKA characterized ESAT-6 & CFP-10, ADCP linked to *Mtb* cell wall IgG, and binding to individual FcRs minimally distinguished the antigens by principal components analyses. These data demonstrate that *Mtb* cell wall and ESAT-6 & CFP-10 elicit different IgG engagement of multiple FcRs to induce cell signaling and activation. Moreover, unique subclass and glycosylation resulting in divergent Fc functions show that antigen specificity has the potential to alter the impact of IgG on *Mtb* pathogenesis.

**Figure 3.**
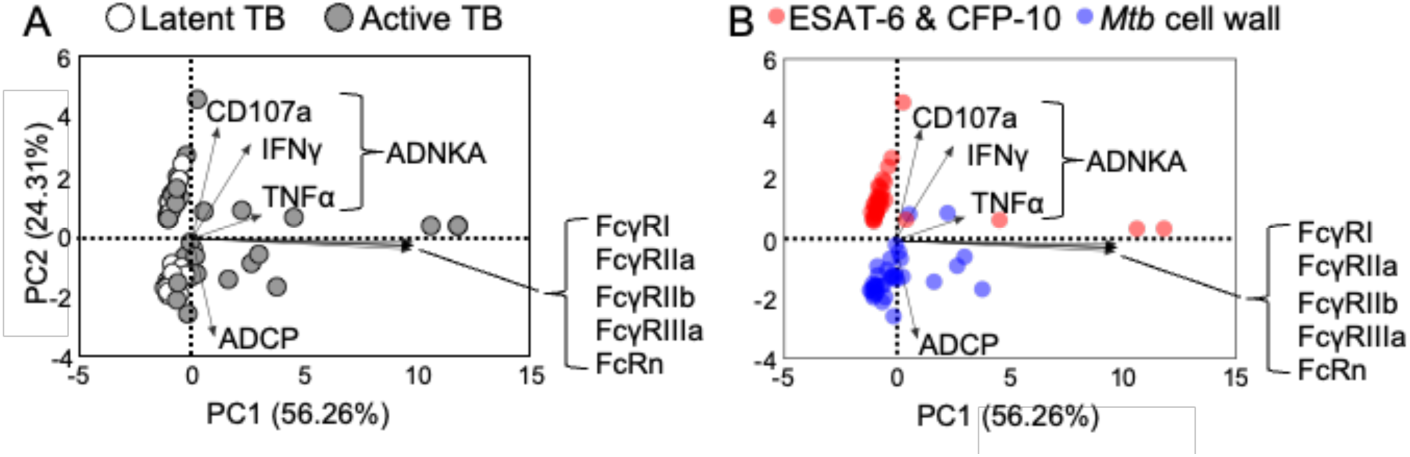
Antibody Fc effector functions diverge more by antigen specificity than latent and active TB. Principal components analysis was performed. Each dot summarizes the linear combination of the Fc receptor binding and Fc effector functions (antibody dependent natural killer cell activation (ADNKA) and antibody dependent cellular phagocytosis (ADCP)) for ESAT-6 & CFP-10 and *Mtb* cell wall IgG in each individual TB patient. Biplots show the distribution of individual TB patient profiles of Fc receptor binding and Fc effector functions to *Mtb* cell wall and ESAT-6 & CFP-10 with respect to **(A)** latent and active TB and **(B)** antigen specificity. The arrows in the loadings show the direction of the Fc features and functions that define the model.

### Latent and active TB IgG differentially impact Mtb in macrophage infection

To evaluate how antigen specificity alters antibody modulation of *Mtb*, we used *in vitro* macrophage models of infection. This quintessential immune cell niche has the capacity to restrict and also permit bacterial replication (*20, 25, 26, 55, 59–64*). Monoclonal and polyclonal antibodies have been shown to mediate differential uptake of extracellular *Mtb* into the macrophage (*25, 26, 61, 62, 65*). In this cohort, to measure the impact of polyclonal IgG on *Mtb* burden resulting from opsonophagocytosis of extracellular bacteria into the macrophage, we used a virulent *Mtb* H37Rv reporter strain (*Mtb*-276) where luminescence correlates with bacterial burden (Supplemental Figure 4A) (*66, 67*). We used primary human monocyte derived macrophages (pMDM) that express FcRs reported to be involved in TB (FcγRI, FcγRIIa, FcγRIIb, FcγRIIIa, and FcRn). We incubated bacteria with IgG from each individual patient and used the opsonized *Mtb* to infect pMDMs at an MOI=1 (Figure 4A). We used a low MOI to mimic the high infectivity and paucibacillary nature of *Mtb* and limit the macrophage toxic effects of non-physiological high loads of bacteria. We quantified *Mtb* growth after infection and burden (Supplemental Figure 4B) and found lower bacterial burden when opsonized by IgG from individuals with latent compared to active TB (Figure 4B). These data demonstrate that polyclonal IgG from individuals across the spectrum of latent and active TB have properties that alter uptake of extracellular *Mtb* into the macrophage and its subsequent growth.

**Figure 4.**
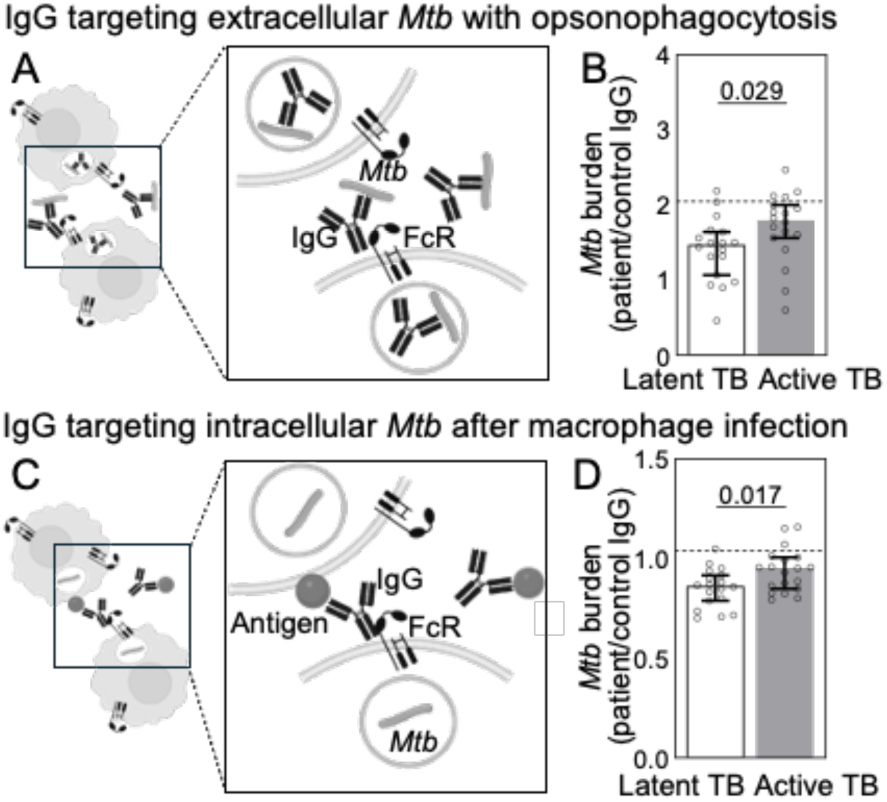
Latent and active TB IgG differentially impact extracellular and intracellular *Mtb* in macrophage infection. **(A)** To test the effect of antibodies on extracellular *Mtb*, IgG isolated from each TB patient was used to opsonize an *Mtb* H37Rv luminescent reporter strain to infect primary human monocyte derived macrophages (MOI=1). **(B)** Each dot represents *Mtb* burden relative to control IgG for each individual TB patient averaged over three different human macrophage donors in independent experiments. Median and 95% CI are shown. The dashed line shows the median of endemic control samples. **(C)** To test the effect of antibodies on intracellular *Mtb*, macrophages were first infected with the virulent *Mtb* H37Rv luminescent reporter strain, extracellular bacteria washed away, *Mtb* infected macrophages treated with IgG, and then **(D)** *Mtb* burden quantitated as in **(B)**. *P*-values are adjusted for sex and age.

While extracellular bacteria are present during initial stages of infection and in advanced disease with active TB, the majority of *Mtb* is intracellular (*59, 60, 68, 69*). We have previously shown that intracellular *Mtb* is differentially impacted by polyclonal IgG pooled from latent and active TB patients (*20*). Here, we enhanced the resolution by evaluating IgG from each individual latent and active TB patient. We infected macrophages first, washed away the extracellular *Mtb*, then treated the *Mtb* infected macrophages with IgG (Figure 4C). Consistent with prior studies, intracellular *Mtb* burden was lower after treatment with IgG from individual latent compared to active TB patients (Figure 4D). Thus, polyclonal IgG from individuals across latent and active TB have properties that impact intracellular *Mtb* replication after infection has occurred. The data from these extracellular (Figure 4B) and intracellular (Figure 4D) assays were then used to understand how antigen specificity affects IgG modulation of *Mtb* macrophage infection.

### Mtb cell wall IgG enhances Mtb burden in opsonophagocytosis of extracellular bacteria

To assess the impact of antigen specificity on opsonophagocytosis of extracellular *Mtb*, we evaluated the relationships between antigen specific IgG properties and *Mtb* burden. Using simple linear regression, we found that *Mtb* burden was positively dependent on *Mtb* cell wall IgG3 and binding to high (FcRn and FcγRI) and low (FcγRIIa and FcγRIIb) affinity FcRs (Figure 5A). These relationships were not observed with control RSV (Supplemental Figure 5). There were no relationships in the context of intracellular bacteria with antibody treatment after macrophage infection (Figure 5B). To assess the impact of IgG glycosylation on *Mtb* burden, we used multiple linear regression to incorporate the combination of glycosylation and subclass. We found that extracellular and not intracellular *Mtb* burden was enhanced by the absence of sialic acid and presence of fucose on IgG (Figure 5C and 5D, Supplemental Table 1 and 2), the glycans that most significantly distinguished *Mtb* cell wall from ESAT-6 & CFP-10 IgG (Figure 2B). As has been reported by others (*25*), these relationships occurred only in latent and not active TB and linked to higher potency of IgG Fc functions (Supplemental Figure 6). These data show that highly potent *Mtb* cell wall IgG enhances bacterial burden by modulating uptake of extracellular *Mtb* into the macrophage.

**Figure 5.**
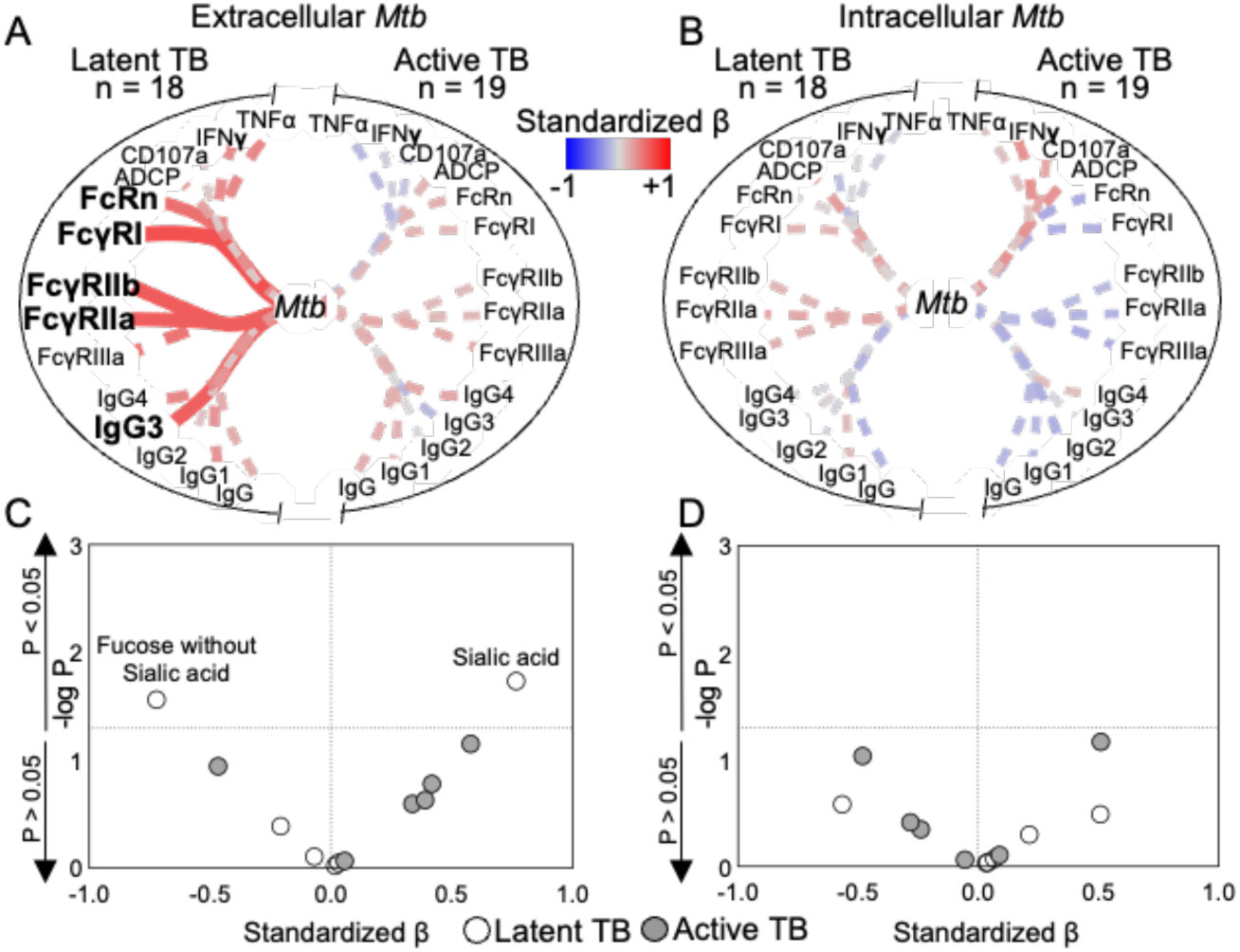
***Mtb* cell wall IgG enhances opsonophagocytosis of extracellular *Mtb.*** The relationships between *Mtb* cell wall IgG features and *Mtb* were determined by simple linear regression in latent (n = 18) and active (n = 19) TB with respect to IgG mediated opsonophagocytosis of extracellular *Mtb* **(A)** and IgG targeting intracellular *Mtb* after macrophage infection **(B)**. Line thickness is inversely proportional to the *P*-value with <0.05 represented by solid lines. The effect of glycans from *Mtb* cell wall IgG on extracellular **(C)** and intracellular **(D)** *Mtb* was determined by multiple linear regression with IgG subclasses (Supplemental Tables 1 and 2). Volcano plots show the strength and direction of the link between *Mtb* cell wall IgG glycans and *Mtb* by standardized β.

### ESAT-6 & CFP-10 IgG inhibits intracellular Mtb

In contrast to *Mtb* cell wall, we observed that ESAT-6 & CFP-10 IgG inhibits *Mtb* (Figure 6A and 6B). More specifically, ESAT-6 & CFP-10 IgG1 negatively linked to intracellular bacterial burden and growth (Figure 6C and 6D) while IgG glycosylation was not (Supplemental Figure 6 and Supplemental Tables 3 and 4). As with *Mtb* cell wall IgG, the observations for ESAT-6 & CFP10 IgG were made in latent and not active TB but functional potency no different (Supplemental Figure 7). These data suggest that ESAT-6 & CFP-10 IgG in latent TB have the capacity to protect.

**Figure 6.**
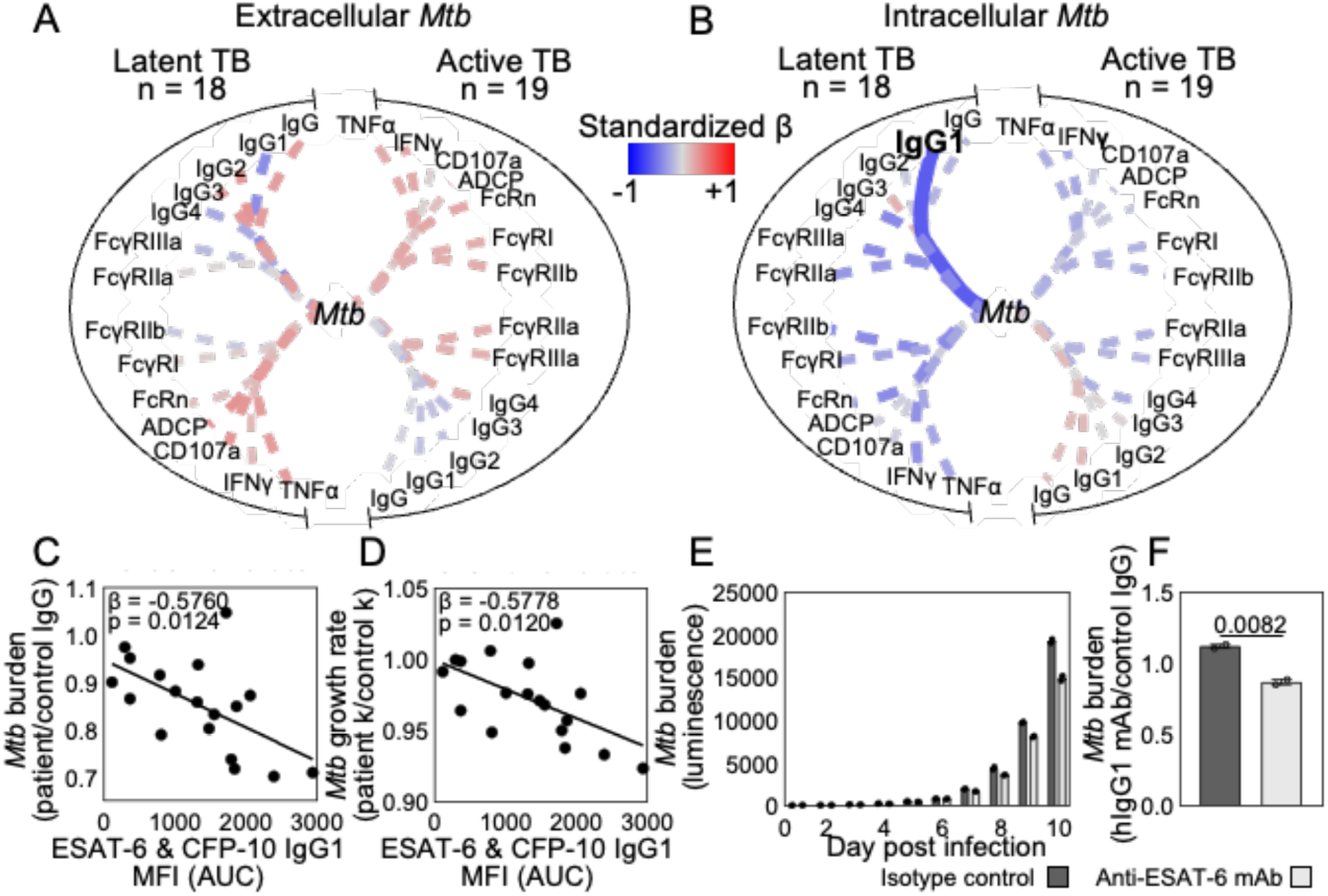
ESAT-6 & CFP-10 IgG inhibits intracellular *Mtb*. The relationships between ESAT-6 & CFP-10 IgG features and *Mtb* were determined by simple linear regression in latent (n = 18) and active (n = 19) TB with respect to IgG mediated opsonophagocytosis of extracellular *Mtb* **(A)** and IgG targeting intracellular *Mtb* after macrophage infection **(B)**. Line thickness is inversely proportional to the *P*-value with <0.05 represented by the solid line. Dot plots show the dependence of intracellular *Mtb* **(C)** burden and **(D)** growth rate on ESAT-6 & CFP-10 IgG1 in latent TB. **(E)** Intracellular *Mtb* burden after treatment with a monoclonal hIgG1 reactive to ESAT-6 & CFP-10 (anti-ESAT-6 mAb) and isotype control are shown for each day post infection by mean and SEM. **(F)** *Mtb* burden relative to control IgG is shown and unpaired t-test used to evaluate significance.

To test the ability of ESAT-6 & CFP-10 IgG1 to inhibit *Mtb*, we cloned a monoclonal human IgG1 (ESAT-6 mAb) that recognizes ESAT-6, CFP-10, and the combination of antigens (Supplemental Figure 8A-C). Treatment of *Mtb* infected macrophages with the ESAT-6 mAb as compared to isotype control led to decreased intracellular *Mtb* burden in a dose-dependent manner (Figure 6E and 6F and Supplemental Figure 8D and 8E). These data show that IgG recognizing the secreted *Mtb* virulence factors ESAT-6 & CFP-10 function by inhibiting intracellular bacteria after infection has been established.

## Discussion

In this study, we use *Mtb* cell wall proteins and the secreted bacterial virulence factors ESAT-6 & CFP-10 to show that antigen specificity alters how antibody Fc effector functions impact *Mtb* pathogenesis. Previous studies using PPD show that different Fc effector functions correlate with protection and disease in latent and active TB. Here, we show that unique antigens within PPD elicit divergent IgG Fc properties in subclass distribution (Figure 1), glycosylation (Figure 2) and Fc effector functions (Figure 3) that change the effect on *Mtb* infection (Figure 4). *Mtb* cell wall IgG enhances disease through opsonophagocytosis of extracellular *Mtb*. This is through IgG3, the subclass with the highest ability to induce Fc receptor binding and effector functions (*70*), and links to FcγRI, FcγRIIa, and FcγRIIb that drive antibody dependent cellular phagocytosis, as well as the high affinity FcRn (Figure 5A). In addition, the presence of sialic acid and absence of fucose on *Mtb* cell wall IgG link to *Mtb* burden (Figure 5C), indicating that glycan modulation of FcR binding as described in the literature is also important (*17, 71–74*). In contrast, ESAT-6 & CFP-10 IgG1 inhibits the intracellular *Mtb* population after infection has occurred (Figure 6B). Its protective capacity is shown in both polyclonal and mAb studies where ESAT-6 & CFP-10 IgG1 alone is sufficient to measurably inhibit bacterial burden (Figure 6E and 6F). These data show that in a single individual TB patient, not all antibodies are equal, and antigen specificity shapes antibody functions that impact *Mtb* pathogenesis.

One possible way that different antigens from the same bacteria could elicit divergent antibody Fc properties could be immune priming. *Mtb* cell wall proteins cross-react with environmental NTMs (*30–34, 42*). In comparison, ESAT-6 & CFP-10 are more specific for *Mtb* such that it is used for current T-cell based diagnostics (*75*). As such, *Mtb* cell wall antibody functions could reflect immunity from exposures to both NTM and *Mtb* whereas ESAT-6 & CFP-10 could represent responses only to *Mtb* (*31*). Further studies that assess the effect of pre-existing NTM immunity on avidity to antigen and Fc properties induced by *Mtb* infection would help evaluate this possibility (*30*).

Antibodies targeting *Mtb* cell wall antigens have been described in the literature to enhance and inhibit *Mtb*. Monoclonal IgG1 targeting the bacterial surface exposed heparin binding hemagglutinin (HBHA) enhance bacterial entry into epithelial cells and subsequent *Mtb* burden through FcRn (*27*). Monoclonal IgG targeting *Mtb* cell wall proteins (PstS1, HspX, and LpqH) (*26, 76, 77*) and monoclonal and polyclonal IgG targeting the bacterial glycan arabinomannan inhibit *Mtb* through Fc dependent and independent mechanisms (*25, 61, 62, 65*). Thus, while the overall mixture of *Mtb* cell wall proteins enhances disease in this study, it is likely that within the cell wall fraction, some individual antigens induce protective and others disease enhancing antibodies.

Antibody dependent enhancement of infection has been described in viral and bacterial infections (*78–81*) including the intracellular pathogens Legionella (*82*) and Leishmania (*83, 84*) in macrophages that involve cross-reactive and FcR mediated mechanisms. Because *Mtb* is also an intracellular pathogen where macrophages represent an important niche, antibody dependent enhancement of infection has been hypothesized to occur in TB. These data show that *Mtb* cell wall polyclonal IgG engage macrophage FcRs to enhance disease.

IgG targeting the secreted virulence factors ESAT-6 & CFP-10 inhibit intracellular bacteria, conferring protection after *Mtb* has established infection (Figure 6B – 6F). Consistent with these findings, the absence and presence of ESAT-6 & CFP-10 in the bacteria is associated with virulence (*85, 86*). Moreover, an anti-ESAT-6 nanobody blocks intracellular *Mtb* replication (*87*). Because the large size of human IgG compared to a nanobody limits cellular penetration, direct neutralization may not be the mechanism of IgG1 mediated bacterial inhibition in an individual TB patient. ESAT-6 & CFP-10 are secreted from *Mtb* and found on the surface of infected host cells (*47, 88, 89*). Thus, it is possible that the secreted nature of these antigens enables immune complex formation, FcR engagement, and induction of effector functions in macrophages to kill intracellular *Mtb*. Our data show that the ability to activate NK cells through FcγRIIIa and induce cellular cytotoxicity highlights ESAT-6 & CFP-10 IgG (Figure 3). A similar process could be occurring with macrophages that express FcγRIIIa and carry out cellular cytotoxicity in addition to FcRs facilitating phagocytosis. mAb Fc modifications to enhance and inhibit cellular cytotoxicity could further evaluate these mechanisms of protection (*90*).

Notably, only latent and not active TB IgG properties impacted *Mtb* burden (Figure 5 and 6). This phenomenon has been reported with arabinomannan specific IgG (*25*) and attributed to higher functional potency. In this study, increased Fc effector functional potency may explain the observations with *Mtb* cell wall, not ESAT-6 & CFP-10 IgG (Supplemental Figure 6). What could explain the phenomenon for ESAT-6 & CFP-10 IgG would be a prozone-like effect from high levels of other *Mtb* reactive IgG in active TB (*91–93*). Additionally, it is possible that differences in antibody functions reflect the continuum of progression and regression in latent TB as compared to the more limited spectrum represented by active TB without antibiotic treatment (*54, 94–96*).

What *Mtb* antigens induce antibody functions that impact the clinical outcomes in TB is unclear but essential to understand if we are to harness antibodies for diagnostics, therapeutics, and vaccines. That both pathogenic and protective IgG exist could explain why some passive serum transfer experiments show inhibition while others enhancement of bacterial burden (*6–9*). *Mtb* cell wall and ESAT-6 & CFP-10 reflect a fraction of the *Mtb* antigen repertoire (*35, 36*). This approach could identify the protective and pathogenic potential for additional bacterial antigens in polyclonal antibody responses and enable us to skew humoral immunity towards beneficial rather than harmful effects. This study represents a starting point with IgG from blood. Extending evaluations to IgA and IgM in pulmonary and peripheral responses will highlight isotype and compartment specific distinctions in mucosal (*30, 76, 97–100*) and systemic immunity critical to understanding the many roles of antibodies in TB.

## Materials and Methods

### Study design

Adults were recruited from the Texas/Mexico border (2006–2010) (*101*). Latent TB (n=18) was defined by a positive interferon γ release assay (IGRA) TSPOT or QuantiFERON with no history of prior TB diagnosis or treatment, and no clinical signs and symptoms of active TB disease. Active TB (n=19) was defined by sputum acid fast bacilli smear and culture in combination with clinical signs and symptoms of disease. Endemic controls (n=8) were defined by negative TSPOT or QuantiFERON with no history, clinical signs and symptoms, or exposure to TB. To limit confounding variables, groups for latent and active TB were matched by age and sex (*54, 102*), tested negative for HIV and type 2 diabetes as defined by WHO criteria (*103–109*), and received <8 days TB treatment (*53, 110*). Written informed consent was obtained from study participants and approved by institutional IRBs.

### Sample Collection

Blood samples were collected by venipuncture in sodium heparin tubes, plasma isolated by centrifugation, aliquoted, stored at −80°C, and heat-inactivated (30 min, 55C) prior to use.

### IgG purification

Polyclonal IgG from patient samples was isolated by negative selection via Melon Gel resin (Thermo Fisher), concentrated by ultra-centrifugal filtration (Millipore), and quantified by ELISA (Mabtech) per manufacturers’ instructions.

### Monoclonal hIgG1 plasmid design and construction

An anti-ESAT-6 VH/k sequence (GenBank: LC189555.1 (*111*)) was synthesized and cloned into a pUC19 vector with a human IgG1 Fc domain as previously described in detail (*90*). Donor and destination plasmids were combined in a single digestion-ligation reaction to generate an expression plasmid encoding the heavy and light chains with BsaI-HF (NEB) and T4 ligase (NEB), transformed into Stellar competent cells (Clontech) and selected by kanamycin.

### Production of monoclonal hIgG1

As previously described (*90*), plasmids were transfected into 293F suspension cells using Polyethylenimine (PEI) (Polysciences). Supernatants were collected 5 days after transfection and IgG was isolated by protein G magnetic beads (16 hours, 4C), eluted using Pierce IgG Elution Buffer (Thermo Fisher), and neutralized with Tris-HCl pH 8.0.

### Cell lines

NK92 cells expressing human FcγRIIIa (CD16.NK-92) (ATCC) were maintained in α-MEM without nucleosides (Thermo Fisher), 2mM L-glutamine (Thermo Fisher), 1.5g/L sodium bicarbonate (Thermo Fisher), 0.02mM folic acid (Alfa Aesar), 0.2mM inositol (MP Biomedicals), 0.1mM β-mercaptoethanol (Thermo Fisher), 100U/mL IL-2 (STEMCELL), 12.5% horse serum (Cytiva), and 12.5% FBS (Gibco), 37C, 5% CO_2_. THP-1 cells (ATCC) were cultured in RPMI-1640 (Sigma-Aldrich), 10% FBS, 2mM L-glutamine, and 10mM HEPES (Thermo Fisher), and 55μM beta-mercaptoethanol, 37C, 5% CO_2_. Freestyle 293F cells (Thermo Fisher) were maintained in a shaking incubator (125RPM, 37C, 8% CO_2_) in Freestyle 293F expression medium (Thermo Fisher).

### Primary human monocyte derived macrophages

Monocytes were isolated from buffy coats obtained from healthy HIV negative adults by CD14 positive selection (Miltenyi) per manufacturer’s instructions and matured by adhesion in RPMI-1640, 10% FBS, 2mM L-glutamine, and 10mM HEPES.

### Mycobacterium tuberculosis H37Rv

A virulent H37Rv (*Mtb*-276) strain expressing luciferase under the P*_hsp60_* promotor was cultured using Middlebrook 7H9 (BD) with 0.05% Tween-80 (Millipore) and Zeocin (20μg/mL) (Invivogen) to log-phase at 37C (*66, 67*), washed with PBS, and passed through a 5µm filter (Millipore) to obtain a single cell suspension prior to infection (MOI=1). Enumeration by colony forming units was performed using serial dilutions on 7H10 medium (BD).

### Antigens

H37Rv purified protein derivative (PPD) (Statens Serum Institute), culture filtrate (BEI), cytosolic proteins (BEI), and cell wall fractions (BEI), as well as ESAT-6 (BEI), CFP-10 (BEI), Ag85A (BEI), Ag85B (BEI) were used as *Mtb* antigens. As controls, respiratory syncytial virus G (BEI) and F (BEI) proteins were used.

### Quantification of antigen specific IgG and subclasses

Customized Luminex assays were used to measure antigen specific IgG and subclass levels as previously described (*73, 112, 113*). Carboxylated microspheres (Bio-Rad, MC100 series) were coupled to protein antigens using an NHS-ester reaction (Thermo Fisher, cat#A32269) following manufacturer’s instructions. Serial dilutions of IgG purified from each individual patient sample (0.18, 0.06, 0.02 ug/mL) was added to antigen-coupled beads (18hours, 4C) and washed. PE-conjugated antibodies detecting total IgG (JDC-10, Southern Biotech), IgG1 (4E3, Southern Biotech), IgG2 (HP6002, Southern Biotech), IgG3 (HP60502, Southern Biotech), and IgG4 (HP6025, Southern Biotech), were added (2 hours, room temperature), washed with PBS 0.05% Tween-20, and re-suspended in PBS to acquire fluorescence intensity on Magpix (Luminex). The relative level of antigen specific antibodies was defined as the area under the curve (AUC) calculated from the serial dilutions for each individual sample.

### Quantification of antigen binding with ESAT-6 mAb

Customized Luminex assays were used to measure antigen binding (*73, 112, 113*). Antigens were coupled to carboxylated microspheres as described above. Serial dilutions of monoclonal hIgG1 (15, 1.5, 0.15 μg/mL) were added to antigen-coupled beads. PE-conjugated anti-human IgG1 was used for detection and fluorescence intensity acquired on Magpix (Luminex) as described above.

### Isolation of antigen specific and total IgG Fc domains

Antigens were biotinylated using EZ-Link Sulfo-NHS-LC-Biotin (Thermo Fisher) per manufacturer’s instructions and coupled to streptavidin beads (NEB). Patient plasma (diluted 1:5 in 0.5M NaCl, 20mM Tris-HCl, 1mM EDTA, pH 7.5) was added to antigen-coupled beads for immunoprecipitation of ESAT-6 & CFP-10 (18 hours, 4C), *Mtb* cell wall (2.5 hours, room temperature), and RSV (18 hours, 4C) specific antibodies. For bulk total IgG, protein G beads (Millipore) were used (2 hours, room temperature). IdeZ (NEB) (1.5 hours, 37C) was used to cleave the Fc domain for subsequent glycan isolation.

### N-linked glycan isolation and quantification

Isolation, labeling, and quantification of N-linked glycosylation are previously described (*73, 113–115*). In brief, isolated Fc domains were denatured (10 minutes, 95C) prior to enzymatic glycan release with PNGaseF (NEB) per manufacturer’s instructions (18 hours, 37C). For antigen specific IgG Fc domains, released glycans were isolated with Agencourt CleanSEQ beads (Beckman Coulter). For total bulk IgG Fc domains, proteins were precipitated in ice-cold ethanol. Glycan-containing supernatants were dried using a CentriVap, then labeled with 8-aminoinopyrene-13,6-trisulfonic acid (APTS) (Thermo Fisher) in 1.2M citric acid and 1M NaBH_3_CN in tetrahydrofuran (Thermo Fisher), and 0.5% NP-40 (NEB) (3 hours, 55C). Excess APTS was removed using Bio-Gel P-2 size exclusion resin (Bio-Rad) (antigen specific glycans) and Agencourt CleanSEQ beads (total bulk glycans). Labeled samples were run with a LIZ 600 DNA ladder (Thermo Fisher) in Hi-Di formamide (Thermo Fisher) on an ABI Gene Analyzer 3500XL and analyzed using GlycanAssure version 1.0 (Thermo Fisher).

### Measurement of antigen specific IgG Fc receptor binding

Customized Luminex assay was used to measure antigen specific IgG FcR binding as previously reported (*73, 112, 113, 116*). As described above, carboxylated microspheres were coupled to protein antigens using an NHS-ester reaction and serial dilutions of IgG purified from each individual patient sample (0.18, 0.06, 0.02 ug/mL) was added to antigen-coupled beads. Recombinant FcRs (FcγRIIIa, FcγRIIa, FcγRIIb, and FcRn) (R&D) were conjugated with phycoerythrin (PE) (Abcam) per manufacturer’s instructions. PE-conjugated FcγRIIIa, FcγRIIa, and FcγRIIb were added at pH 7.4; PE-conjugated FcRn at pH 6.0 (*117*) (2 hours, room temperature). FcγRI recombinant protein (R&D) was added to IgG coated beads and then incubated with mouse anti-human FcγRI (10.1, Santa Cruz) (1 hour, room temperature) followed by PE-conjugated goat anti-mouse (Southern Biotech) (1 hour, room temperature) for detection. Fluorescence intensity was acquired on a Magpix instrument (Luminex), and AUC calculated as described above.

### Antibody dependent cellular phagocytosis

The THP-1 phagocytosis assay of antigen-coated beads is previously described (*73, 113, 118*). *Mtb* cell wall, ESAT-6 & CFP-10, or RSV antigen mix were biotinylated with EZ-Link Sulfo-NHS-LC-Biotin following manufacturer’s instructions and coupled to FluoSpheres NeutrAvidin beads (Molecular Probes) (16 hours, 4C). Antigen-coupled beads were incubated with 100μg/mL polyclonal IgG purified from each patient and then added to THP-1 cells (1x10^5 per well) (37°C, 16 hours). After fixation with 4% PFA, bead uptake was measured by flow cytometry on a BD-LSR Fortessa and analyzed by FlowJo v10. Phagocytic scores were calculated as the integrated median fluorescence intensity (MFI) (% bead-positive frequency × MFI/10,000) (*119*).

### Antibody dependent natural killer cell activation

Antibody dependent NK cell activation is previously described (*73, 113, 120*). ELISA plates were coated with antigen (300ng/well) (18 hours, 4C), washed 3 times with PBS, blocked with 5% BSA (18 hours, 4C), and then washed 3 times. Purified polyclonal IgG from each patient (100μg/mL) was added (2 hours, 37C), followed by CD16a.NK-92 cells (5 × 10^4^ cells/well) with brefeldin A (Biolegend), Golgi Stop (BD Biosciences) and anti-CD107a (H4A3, Biolegend) (5 hours, 37C). Cells were stained with anti-CD56 (5.1H11, Biolegend) and anti-CD16 (3G8, Biolegend) and fixed with 4% PFA. Intracellular cytokine staining to detect IFNγ (B27, Biolegend) and TNFα (Mab11, BD Biosciences) was performed in permeabilization buffer (Biolegend). Markers were measured using a BD LSR Fortessa and analyzed by FlowJo as described above.

### Macrophage Mtb infections

To test the impact of antibodies on the uptake of extracellular *Mtb* into the macrophage and its subsequent replication, purified IgG from each individual patient was incubated with log-phase *Mtb* (4 hours, 37C). The IgG (100ug/mL) opsonized *Mtb* were used to infect primary human monocyte derived macrophages (5x10^4 cells/well) at an MOI=1.

To test the impact of antibodies on intracellular *Mtb* replication, macrophages were first infected with *Mtb* (MOI=1) (14 hours, 37C). Extracellular *Mtb* was then washed off prior to the addition of IgG from each individual patient (100μg/mL).

Luminescence was measured using a BioTek Synergy Neo2 Hybrid Multimode Reader plate reader every 24 hours until *Mtb* growth reached stationary phase. Each patient sample was tested in duplicate in three independent experiments using macrophage derived from three different healthy HIV negative donors.

### Statistics

Data are presented as median with 95% confidence intervals (Figure 1B-C, 4B,4D, Supplemental Figure 1, 3A). Data was analyzed by Mann – Whitney test (Table), Chi-square test (Table), logistic regression (Figure 1A), multiple linear regression to adjust for age and sex (Figure 1B-C, 4B, 4D, Supplemental Figure 1, 3A, 7) and to incorporate multiple antibody features (Figure 5C,5D, Supplemental Tables 1-4, Supplemental Figure 6), Friedman test with adjustment for FDR using a two-stage step-up method of Benjamini, Krieger, and Yekutieli with Q=1% (Figure 2B, and Supplemental Figure 2D), principal components analysis (Figure 2C, 2D, 3A-B, Supplemental Figure 2E), hierarchical clustering (Supplemental Figure 2F), simple linear regression (Figure 5A, 5B, 6A, 6B, Supplemental Figure 5A-B), Pearson correlation (Supplemental Figure 4A), and unpaired t-test (Figure 6F) using STATA v16, Graphpad Prism10, and JMP17.2.0. Figures were generated using Graphpad Prism10, R using the ggplot package, Cytoscape v3.9.1, JMP17.2.0, and Biorender.

## Author contributions

JM and PL designed and conducted experiments, acquired data, analyzed data, and wrote the manuscript. SB conducted experiments and acquired data. GA and JR provided reagents. BG conducted experiments, acquired data, analyzed data, provided reagents, and wrote the manuscript. BR designed the research study, provided reagents, and wrote the manuscript. LL designed the research study, conducted experiments, acquired data, analyzed data, and wrote the manuscript.

## Acknowledgements

We thank Gabrielle Lessen for her assistance with the in vitro *Mtb* macrophage infection experiments and manuscript editing. We thank Michael Shiloh for helpful comments. We thank study participants and clinical staff from the Secretaria de Salud de Tamaulipas and the Hidalgo Department of State and Health Services for field logistics support. This work is supported by UT Southwestern Disease Oriented Scholars Award (LLL), NIH NIAID 5R01AI158858 (LLL), NIH NIAID R21AI144541 (BIR), and NIH NIAID 5T32AI005284-44 (JRM).

---

**Supplemental Figure 1.**
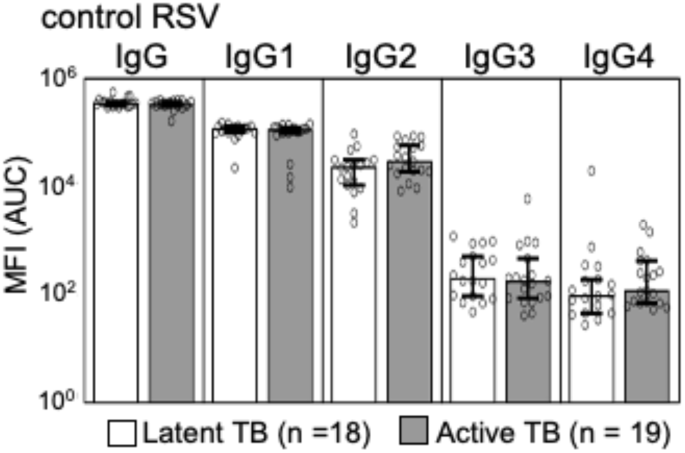
No difference in RSV specific IgG between latent and active TB. Bar graphs show the median and 95% CI of IgG reactive to RSV. No *P-*values adjusted for sex and age were ≤ 0.05.

**Supplemental Figure 2.**
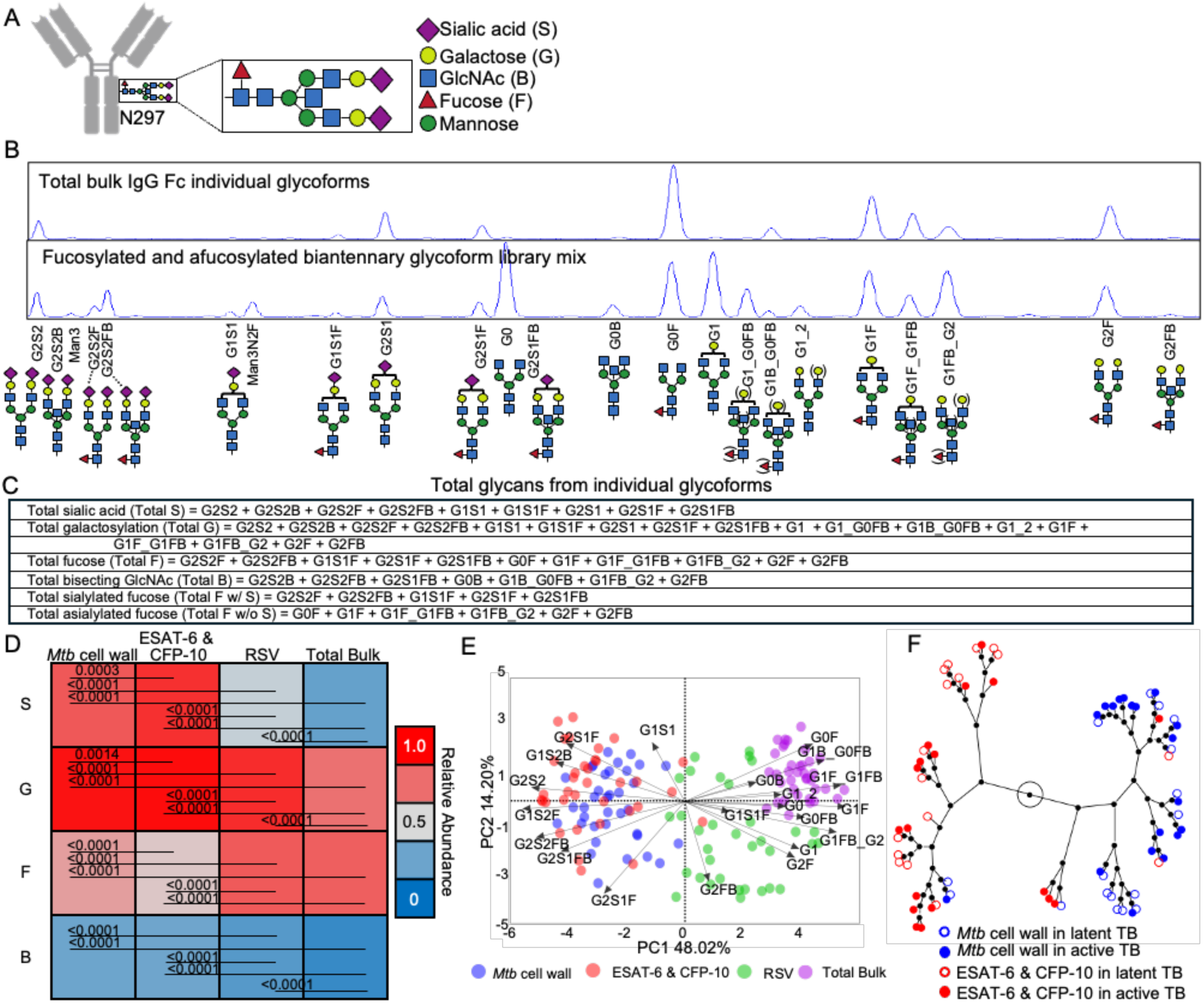
Antigen specificity impacts IgG Fc domain N-glycosylation. **(A)** The conserved N297-linked complex biantennary glycan structure of the IgG Fc domain is shown. **(B)** Individual glycoforms on polyclonal IgG are quantified by capillary electrophoresis using standards from libraries. **(C)** Total glycans summarize individual glycoforms. **(D)** Heatmap shows the relative abundance of total glycans by antigen specificity. Comparisons were made by Wilcoxon matched-pairs signed rank tests and adjusted for multiple comparisons. **(E)** The biplot shows the distribution of individual TB patient glycoform patterns for *Mtb* cell wall, ESAT-6 & CFP-10, RSV, and total bulk IgG. The arrows in the loadings show the direction of the individual glycoforms that define the model. **(F)** The constellation plot shows the relationships between the glycan patterns for ESAT6 & CFP10 and *Mtb* cell wall IgG in each individual TB patient as determined by hierarchical clustering.

**Supplemental Figure 3.**
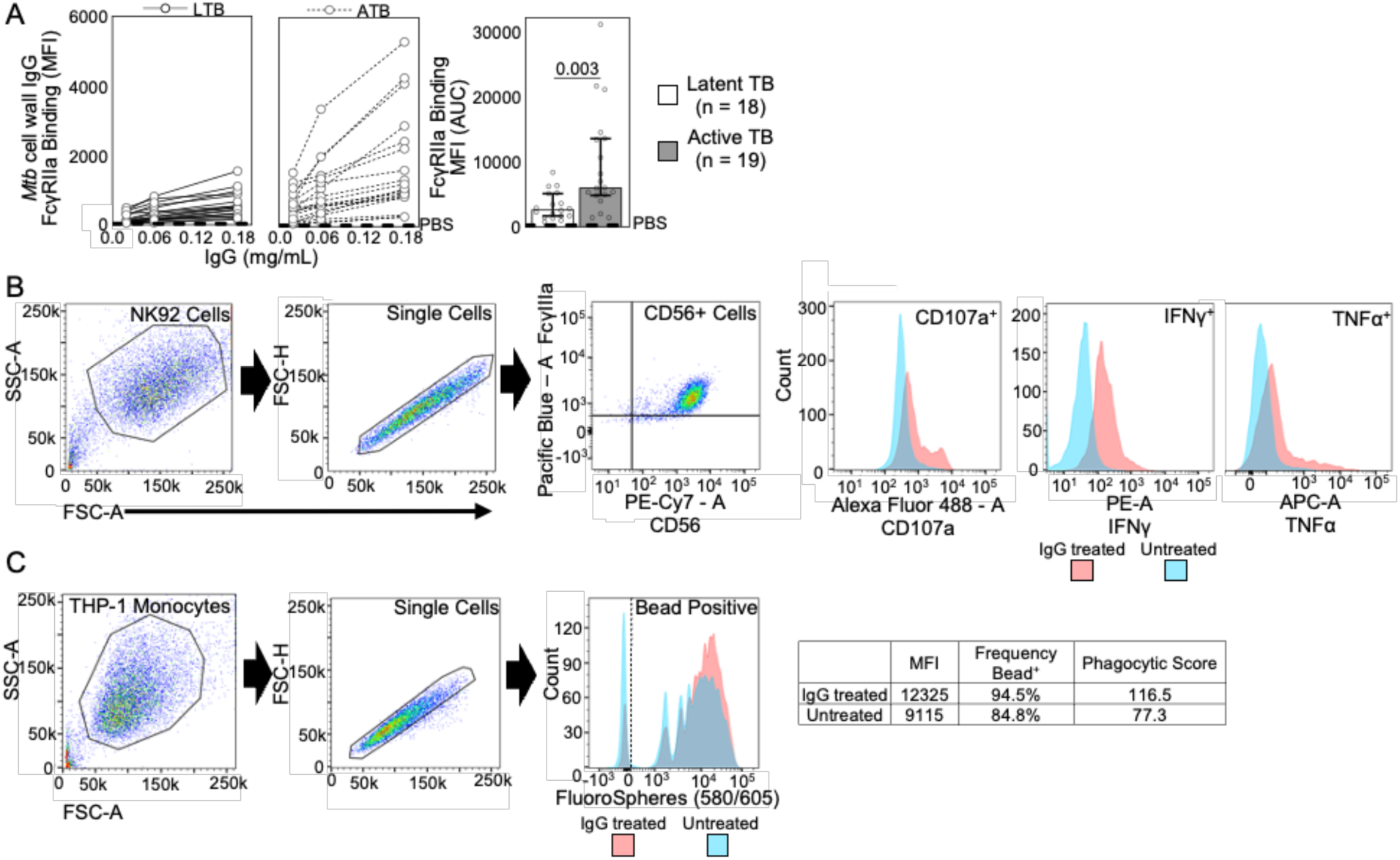
Binding to Fc receptors, antibody dependent natural killer cell activation (ADNKA), and antibody dependent cellular phagocytosis (ADCP) was determined for each individual TB patient sample for *Mtb* cell wall and ESAT-6 & CFP-10. **(A)** Customized Luminex assays were used to measure antigen specific IgG-FcγR binding for each TB patient sample across three dilutions. *Mtb* cell wall IgG binding to FcγIIa plots are shown as examples. Area under the curve (AUC) of the MFI are determined for each individual TB patient and depicted in bar graphs with the median and 95% CI and significance adjusted for sex and age. **(B)** Antigen specific antibody dependent NK cell activation (ADNKA) was determined for each individual patient sample using the NK92 expressing FcγRIIIa cell line and antigen adsorbed ELISA plates. CD107a, IFNg, and TNFa on CD56+ cells were used to measure activation. **(C)** Antibody dependent cellular phagocytosis (ADCP) was quantified for each individual patient sample using THP-1 monocytes and antigen adsorbed fluorescent beads. The phagocytic score was calculated as the integrated MFI (%frequency x MFI/10,000).

**Supplemental Figure 4.**
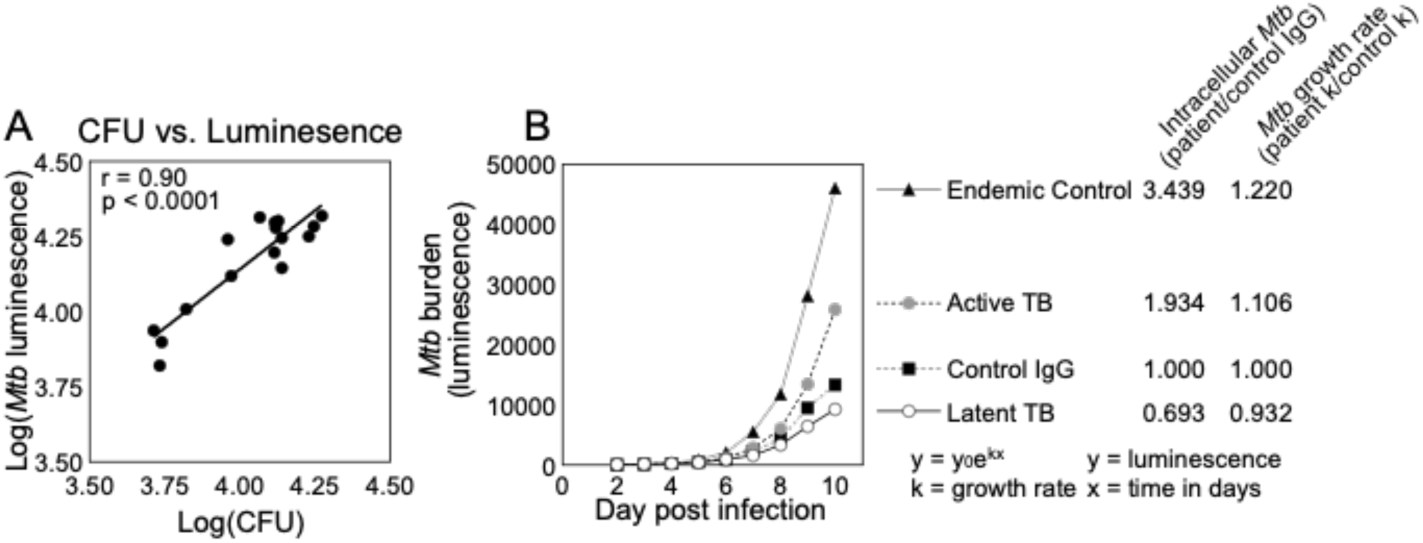
The virulent *Mtb* H37Rv luminescent reporter strain captures differences in bacterial growth and burden between treatment with latent and active TB IgG. **(A)** Pearson correlation was used to link luminescence and CFU. **(B)** Daily luminescence measurements of *Mtb* infected macrophages enable the determination of bacterial burden and growth rate by the exponential growth model.

**Supplemental Figure 5.**
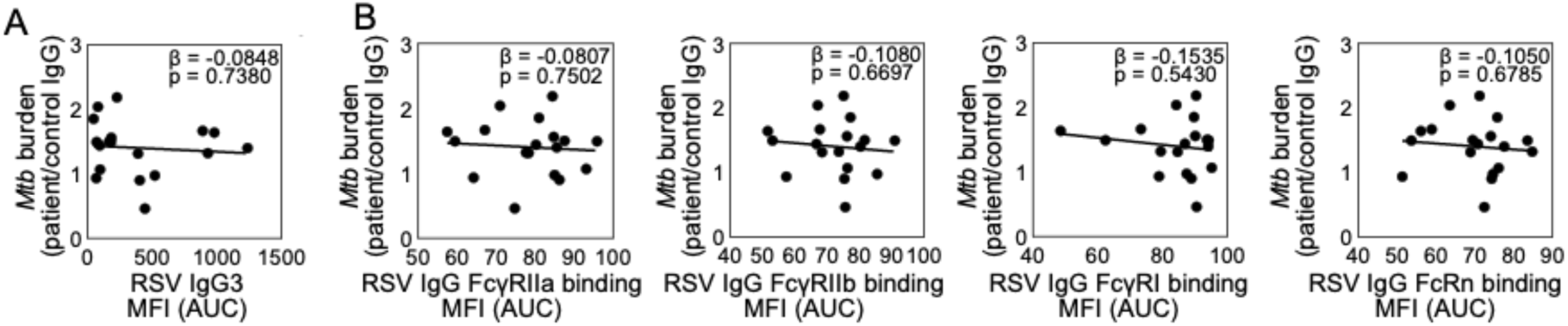
RSV IgG3 and Fc receptor binding do not relate to extracellular *Mtb*. The absence of significant relationships between *Mtb* burden after IgG mediated opsonophagocytosis of extracellular bacteria and **(A)** RSV IgG3 and **(B)** RSV IgG Fc receptor binding was determined by linear regression.

**Supplemental Figure 6.**
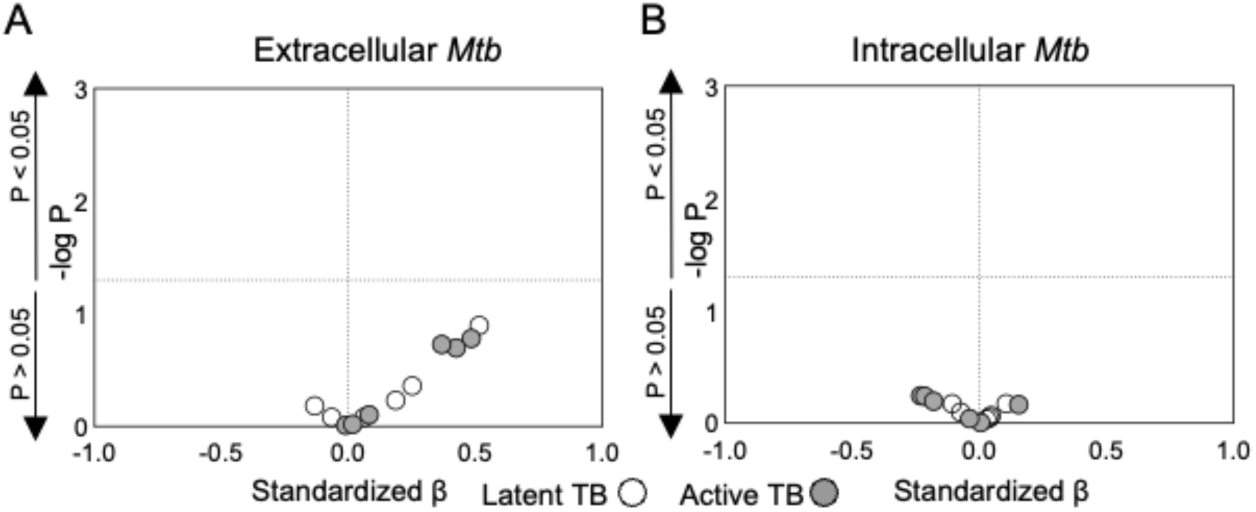
ESAT-6 & CFP-10 glycans do not relate to *Mtb* burden. The effect of glycans from ESAT-6 & CFP-10 IgG on extracellular (A) and intracellular (B) Mtb was determined by multiple linear regression with IgG subclasses (Supplemental Tables 3 and 4). Volcano plots show the strength and direction of the link between ESAT-6 & CFP-10 IgG glycans and *Mtb* by standardized β.

**Supplemental Figure 7.**
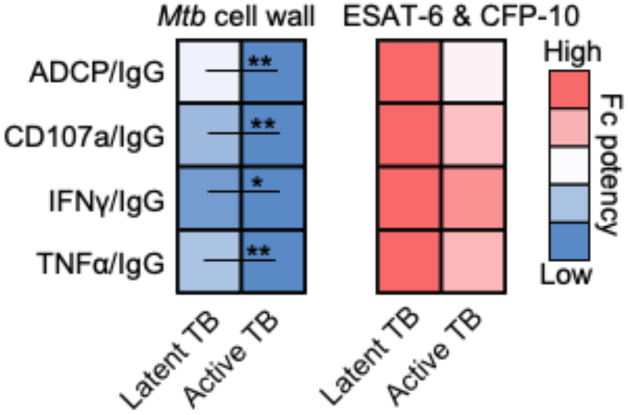
M*t*b cell wall, not ESAT-6 & CFP-10, IgG Fc potency is higher in latent compared to active TB. The heatmap shows the amount of antigen specific Fc effector functions relative to IgG titers with significance adjusted for sex and age. \**P* ≤ 0.05 and \*\**P* ≤ 0.01.

**Supplemental Figure 8.**
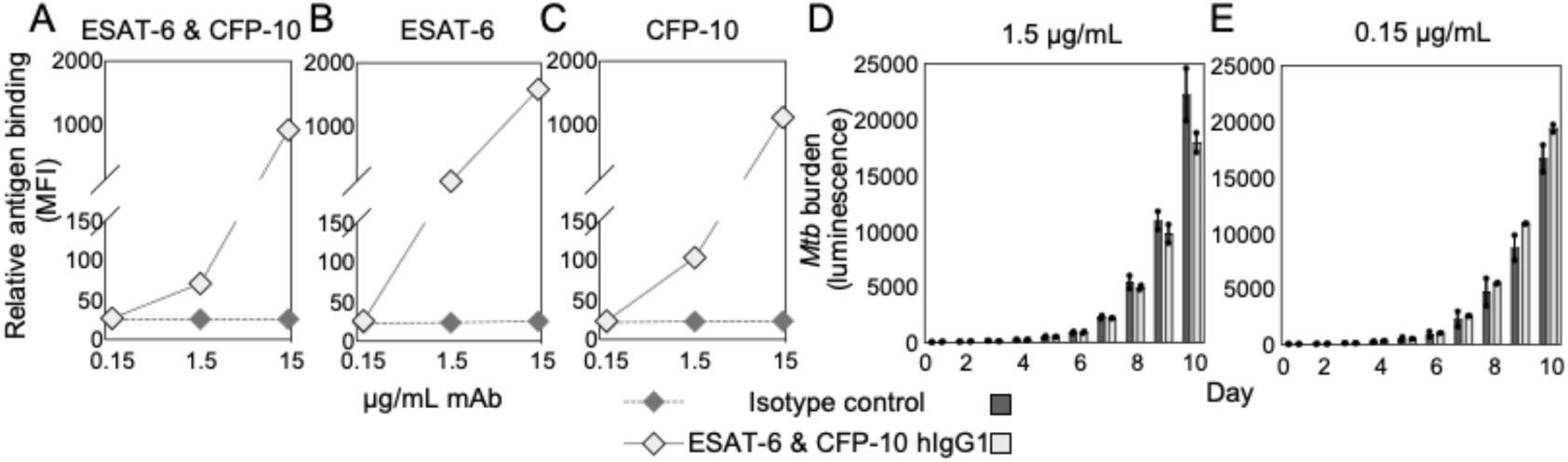
A human IgG1 mAb binds ESAT-6 & CFP-10 and does not inhibit intracellular *Mtb* at low doses. Relative binding of mAb to ESAT-6 and isotype control CR3022 (anti-SARS-CoV-2 RBD hIgG1) to **(A)** ESAT-6 & CFP-10, **(B)** ESAT-6, and **(C)** CFP-10 are shown. Treatment of *Mtb* infected macrophages with **(D)** 1.5 or **(E)** 0.15μg/mL mAb did not significantly impact *Mtb* burden compared to isotype control as determined by unpaired t-test. Mean and SEM are shown.

**Table:** Cohort characteristics.

|  | Latent TB | Active TB | Significance | Endemic Controls |
| --- | --- | --- | --- | --- |
| Total number of individuals | 18 | 19 |  | 8 |
| Age in years (range [median]) | 23 – 79 (40) | 18 – 82 (38) | $P = 0.5730^A$ | 30 – 65 (47) |
| Sex (no. [%]) | | | $P = 0.6185^B$ | |
| Female | 10 (56%) | 9 (47%) |  | 5 (63%) |
| Male | 8 (44%) | 10 (53%) |  | 3 (38%) |
| BCG vaccination | 15 (83%) | 16 (84%) | $P = 0.9423^C$ | 7 (88%) |

<sup>A</sup>Mann-Whitney U test was used to evaluate ages and <sup>B,C</sup>Chi-square test for sex and BCG vaccination.

**Supplemental Table 1.**
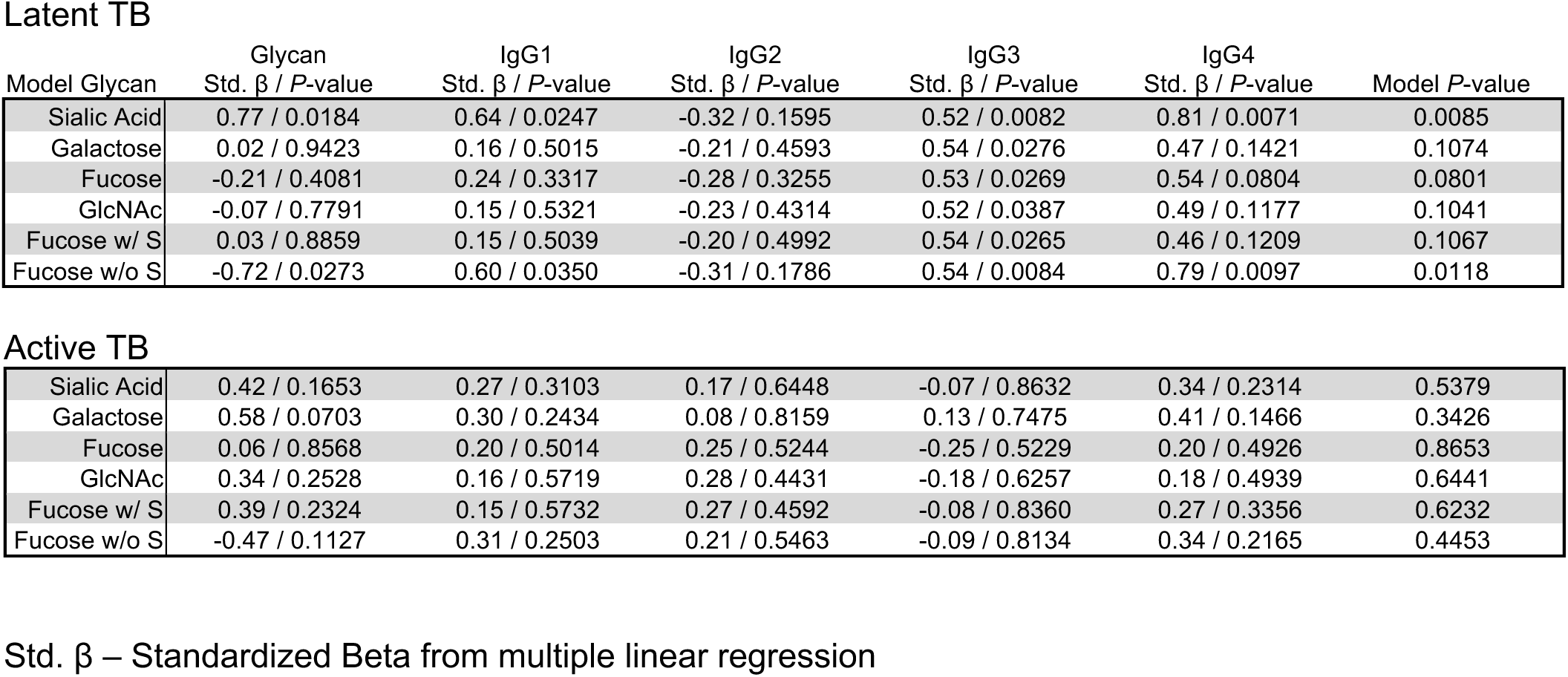
Models evaluating the impact of *Mtb* cell wall IgG glycans and subclasses on extracellular *Mtb*.

**Latent TB**
| Model Glycan | Glycan<br>Std. $\beta$ / *P*-value | IgG1<br>Std. $\beta$ / *P*-value | IgG2<br>Std. $\beta$ / *P*-value | IgG3<br>Std. $\beta$ / *P*-value | IgG4<br>Std. $\beta$ / *P*-value | Model *P*-value |
| --- | --- | --- | --- | --- | --- | --- |
| Sialic Acid | 0.77 / 0.0184 | 0.64 / 0.0247 | -0.32 / 0.1595 | 0.52 / 0.0082 | 0.81 / 0.0071 | 0.0085 |
| Galactose | 0.02 / 0.9423 | 0.16 / 0.5015 | -0.21 / 0.4593 | 0.54 / 0.0276 | 0.47 / 0.1421 | 0.1074 |
| Fucose | -0.21 / 0.4081 | 0.24 / 0.3317 | -0.28 / 0.3255 | 0.53 / 0.0269 | 0.54 / 0.0804 | 0.0801 |
| GlcNAc | -0.07 / 0.7791 | 0.15 / 0.5321 | -0.23 / 0.4314 | 0.52 / 0.0387 | 0.49 / 0.1177 | 0.1041 |
| Fucose w/ S | 0.03 / 0.8859 | 0.15 / 0.5039 | -0.20 / 0.4992 | 0.54 / 0.0265 | 0.46 / 0.1209 | 0.1067 |
| Fucose w/o S | -0.72 / 0.0273 | 0.60 / 0.0350 | -0.31 / 0.1786 | 0.54 / 0.0084 | 0.79 / 0.0097 | 0.0118 |
| **Active TB** |  |  |  |  |  |  |
| Sialic Acid | 0.42 / 0.1653 | 0.27 / 0.3103 | 0.17 / 0.6448 | -0.07 / 0.8632 | 0.34 / 0.2314 | 0.5379 |
| Galactose | 0.58 / 0.0703 | 0.30 / 0.2434 | 0.08 / 0.8159 | 0.13 / 0.7475 | 0.41 / 0.1466 | 0.3426 |
| Fucose | 0.06 / 0.8568 | 0.20 / 0.5014 | 0.25 / 0.5244 | -0.25 / 0.5229 | 0.20 / 0.4926 | 0.8653 |
| GlcNAc | 0.34 / 0.2528 | 0.16 / 0.5719 | 0.28 / 0.4431 | -0.18 / 0.6257 | 0.18 / 0.4939 | 0.6441 |
| Fucose w/ S | 0.39 / 0.2324 | 0.15 / 0.5732 | 0.27 / 0.4592 | -0.08 / 0.8360 | 0.27 / 0.3356 | 0.6232 |
| Fucose w/o S | -0.47 / 0.1127 | 0.31 / 0.2503 | 0.21 / 0.5463 | -0.09 / 0.8134 | 0.34 / 0.2165 | 0.4453 |

Std. $\beta$ – Standardized Beta from multiple linear regression

**Supplemental Table 2.**
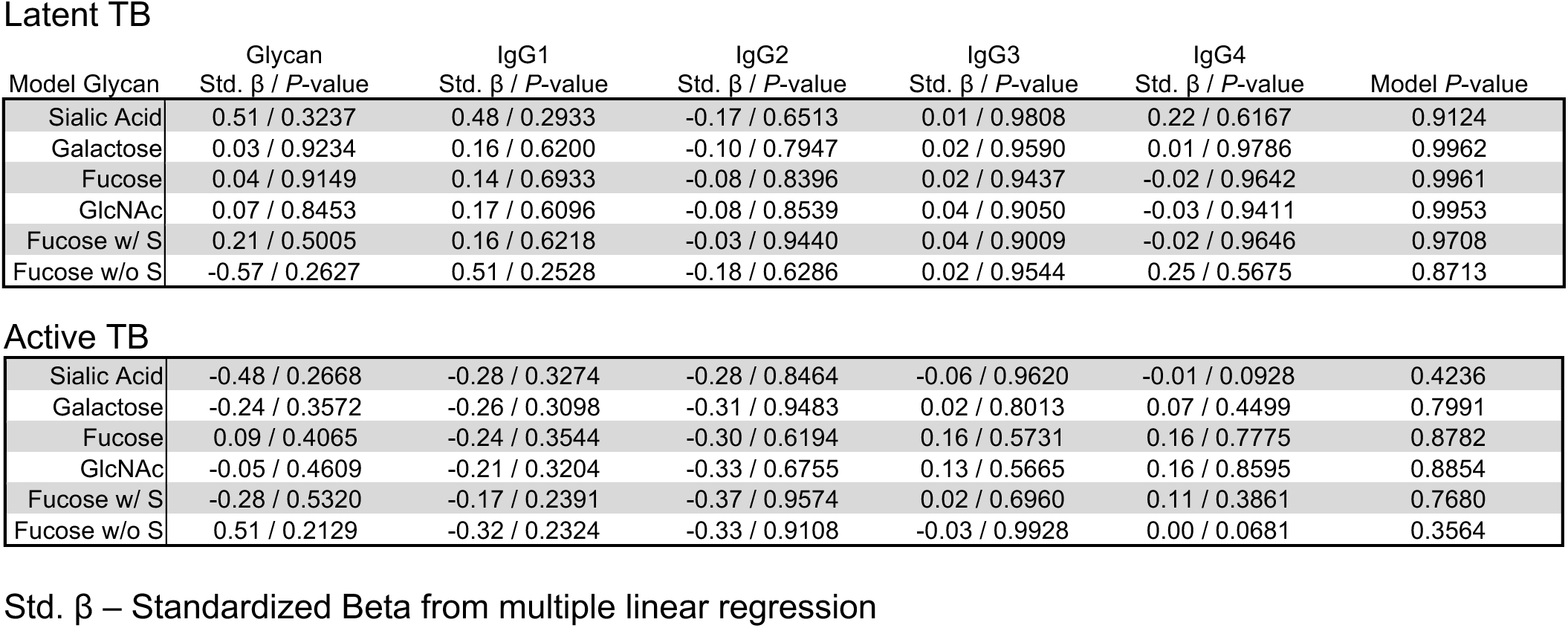
Models evaluating the impact of *Mtb* cell wall IgG glycans and subclasses on intracellular *Mtb*.

**Latent TB**
| Model Glycan | Glycan<br>Std. $\beta$ / *P*-value | IgG1<br>Std. $\beta$ / *P*-value | IgG2<br>Std. $\beta$ / *P*-value | IgG3<br>Std. $\beta$ / *P*-value | IgG4<br>Std. $\beta$ / *P*-value | Model *P*-value |
| --- | --- | --- | --- | --- | --- | --- |
| Sialic Acid | 0.51 / 0.3237 | 0.48 / 0.2933 | -0.17 / 0.6513 | 0.01 / 0.9808 | 0.22 / 0.6167 | 0.9124 |
| Galactose | 0.03 / 0.9234 | 0.16 / 0.6200 | -0.10 / 0.7947 | 0.02 / 0.9590 | 0.01 / 0.9786 | 0.9962 |
| Fucose | 0.04 / 0.9149 | 0.14 / 0.6933 | -0.08 / 0.8396 | 0.02 / 0.9437 | -0.02 / 0.9642 | 0.9961 |
| GlcNAc | 0.07 / 0.8453 | 0.17 / 0.6096 | -0.08 / 0.8539 | 0.04 / 0.9050 | -0.03 / 0.9411 | 0.9953 |
| Fucose w/ S | 0.21 / 0.5005 | 0.16 / 0.6218 | -0.03 / 0.9440 | 0.04 / 0.9009 | -0.02 / 0.9646 | 0.9708 |
| Fucose w/o S | -0.57 / 0.2627 | 0.51 / 0.2528 | -0.18 / 0.6286 | 0.02 / 0.9544 | 0.25 / 0.5675 | 0.8713 |
| **Active TB** |  |  |  |  |  |  |
| Sialic Acid | -0.48 / 0.2668 | -0.28 / 0.3274 | -0.28 / 0.8464 | -0.06 / 0.9620 | -0.01 / 0.0928 | 0.4236 |
| Galactose | -0.24 / 0.3572 | -0.26 / 0.3098 | -0.31 / 0.9483 | 0.02 / 0.8013 | 0.07 / 0.4499 | 0.7991 |
| Fucose | 0.09 / 0.4065 | -0.24 / 0.3544 | -0.30 / 0.6194 | 0.16 / 0.5731 | 0.16 / 0.7775 | 0.8782 |
| GlcNAc | -0.05 / 0.4609 | -0.21 / 0.3204 | -0.33 / 0.6755 | 0.13 / 0.5665 | 0.16 / 0.8595 | 0.8854 |
| Fucose w/ S | -0.28 / 0.5320 | -0.17 / 0.2391 | -0.37 / 0.9574 | 0.02 / 0.6960 | 0.11 / 0.3861 | 0.7680 |
| Fucose w/o S | 0.51 / 0.2129 | -0.32 / 0.2324 | -0.33 / 0.9108 | -0.03 / 0.9928 | 0.00 / 0.0681 | 0.3564 |

Std. $\beta$ – Standardized Beta from multiple linear regression

**Supplemental Table 3.**
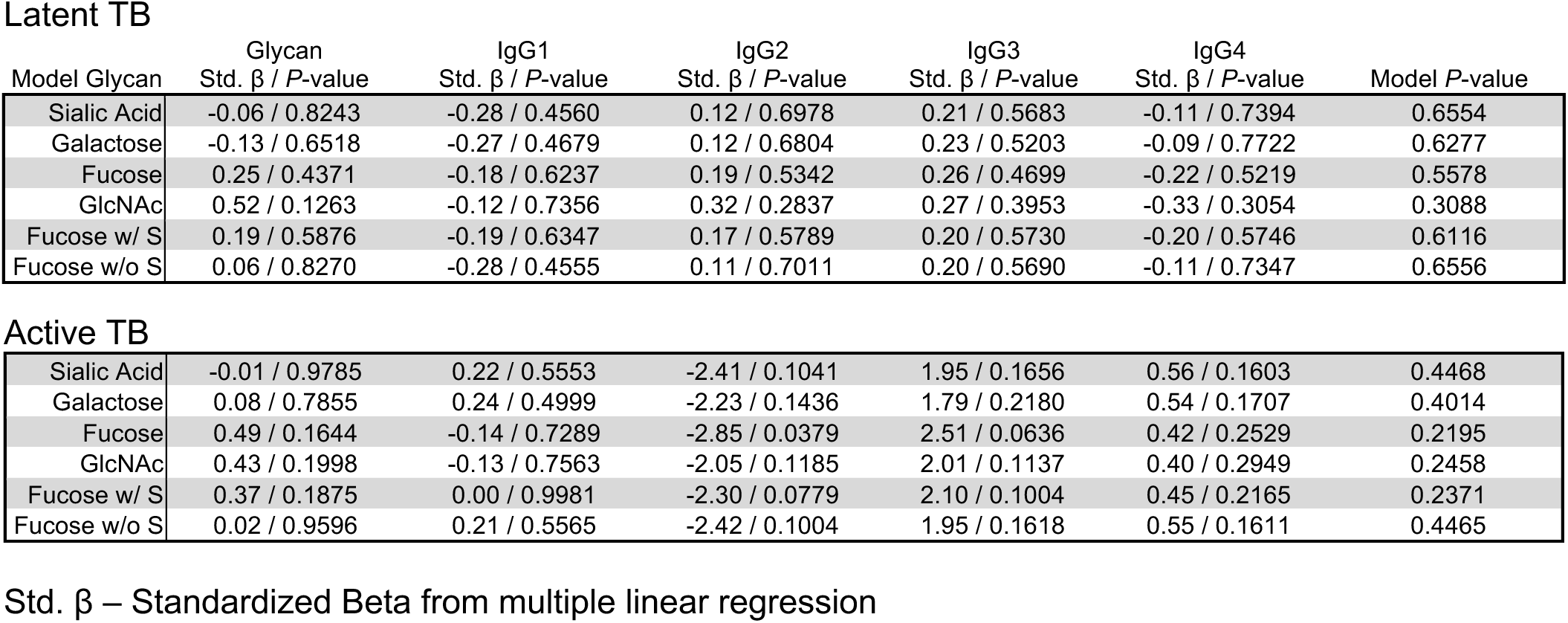
Models evaluating the impact of ESAT-6 & CFP-10 IgG glycans and subclasses on extracellular *Mtb*.

**Latent TB**
| Model Glycan | Glycan<br>Std. $\beta$ / *P*-value | IgG1<br>Std. $\beta$ / *P*-value | IgG2<br>Std. $\beta$ / *P*-value | IgG3<br>Std. $\beta$ / *P*-value | IgG4<br>Std. $\beta$ / *P*-value | Model *P*-value |
| --- | --- | --- | --- | --- | --- | --- |
| Sialic Acid | -0.06 / 0.8243 | -0.28 / 0.4560 | 0.12 / 0.6978 | 0.21 / 0.5683 | -0.11 / 0.7394 | 0.6554 |
| Galactose | -0.13 / 0.6518 | -0.27 / 0.4679 | 0.12 / 0.6804 | 0.23 / 0.5203 | -0.09 / 0.7722 | 0.6277 |
| Fucose | 0.25 / 0.4371 | -0.18 / 0.6237 | 0.19 / 0.5342 | 0.26 / 0.4699 | -0.22 / 0.5219 | 0.5578 |
| GlcNAc | 0.52 / 0.1263 | -0.12 / 0.7356 | 0.32 / 0.2837 | 0.27 / 0.3953 | -0.33 / 0.3054 | 0.3088 |
| Fucose w/ S | 0.19 / 0.5876 | -0.19 / 0.6347 | 0.17 / 0.5789 | 0.20 / 0.5730 | -0.20 / 0.5746 | 0.6116 |
| Fucose w/o S | 0.06 / 0.8270 | -0.28 / 0.4555 | 0.11 / 0.7011 | 0.20 / 0.5690 | -0.11 / 0.7347 | 0.6556 |
| **Active TB** |  |  |  |  |  |  |
| Sialic Acid | -0.01 / 0.9785 | 0.22 / 0.5553 | -2.41 / 0.1041 | 1.95 / 0.1656 | 0.56 / 0.1603 | 0.4468 |
| Galactose | 0.08 / 0.7855 | 0.24 / 0.4999 | -2.23 / 0.1436 | 1.79 / 0.2180 | 0.54 / 0.1707 | 0.4014 |
| Fucose | 0.49 / 0.1644 | -0.14 / 0.7289 | -2.85 / 0.0379 | 2.51 / 0.0636 | 0.42 / 0.2529 | 0.2195 |
| GlcNAc | 0.43 / 0.1998 | -0.13 / 0.7563 | -2.05 / 0.1185 | 2.01 / 0.1137 | 0.40 / 0.2949 | 0.2458 |
| Fucose w/ S | 0.37 / 0.1875 | 0.00 / 0.9981 | -2.30 / 0.0779 | 2.10 / 0.1004 | 0.45 / 0.2165 | 0.2371 |
| Fucose w/o S | 0.02 / 0.9596 | 0.21 / 0.5565 | -2.42 / 0.1004 | 1.95 / 0.1618 | 0.55 / 0.1611 | 0.4465 |

Std. $\beta$ – Standardized Beta from multiple linear regression

**Supplemental Table 4.**
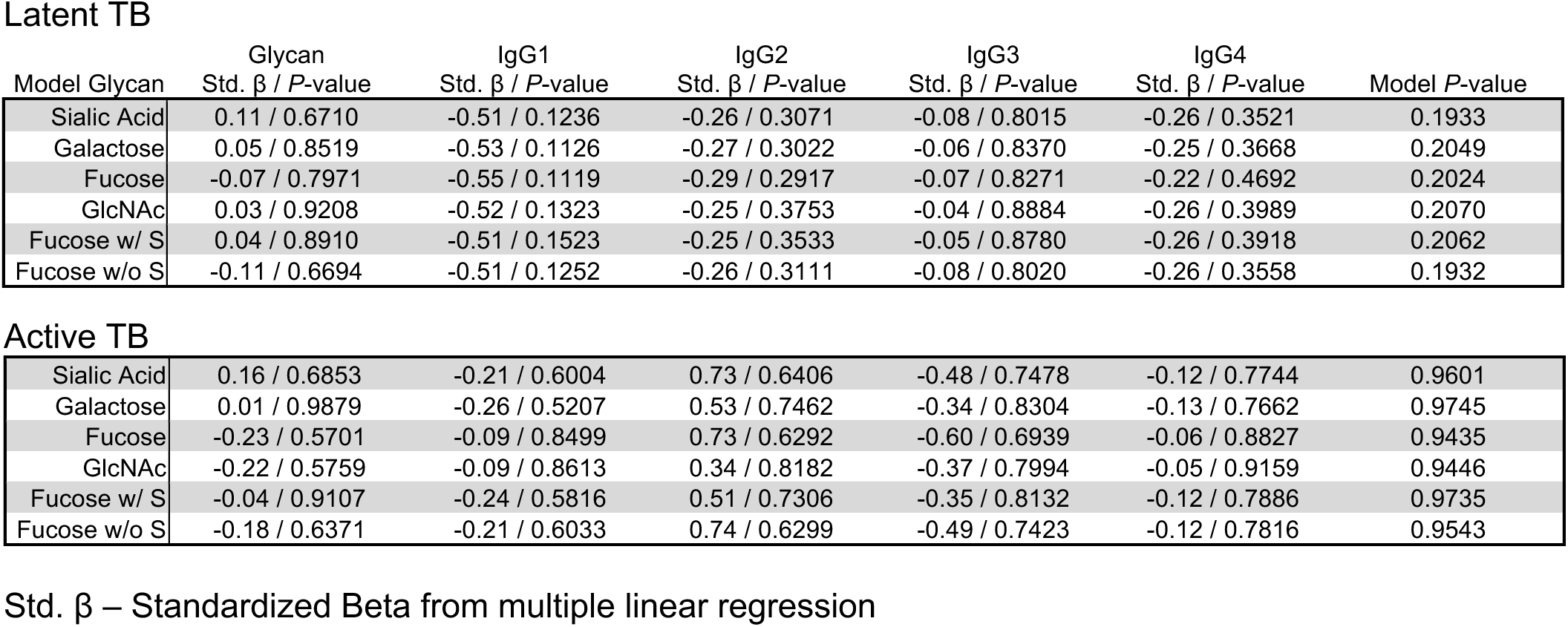
Models evaluating the impact of ESAT-6 & CFP-10 IgG glycans and subclasses on intracellular *Mtb*.

**Latent TB**
| Model Glycan | Glycan<br>Std. $\beta$ / *P*-value | IgG1<br>Std. $\beta$ / *P*-value | IgG2<br>Std. $\beta$ / *P*-value | IgG3<br>Std. $\beta$ / *P*-value | IgG4<br>Std. $\beta$ / *P*-value | Model *P*-value |
| --- | --- | --- | --- | --- | --- | --- |
| Sialic Acid | 0.11 / 0.6710 | -0.51 / 0.1236 | -0.26 / 0.3071 | -0.08 / 0.8015 | -0.26 / 0.3521 | 0.1933 |
| Galactose | 0.05 / 0.8519 | -0.53 / 0.1126 | -0.27 / 0.3022 | -0.06 / 0.8370 | -0.25 / 0.3668 | 0.2049 |
| Fucose | -0.07 / 0.7971 | -0.55 / 0.1119 | -0.29 / 0.2917 | -0.07 / 0.8271 | -0.22 / 0.4692 | 0.2024 |
| GlcNAc | 0.03 / 0.9208 | -0.52 / 0.1323 | -0.25 / 0.3753 | -0.04 / 0.8884 | -0.26 / 0.3989 | 0.2070 |
| Fucose w/ S | 0.04 / 0.8910 | -0.51 / 0.1523 | -0.25 / 0.3533 | -0.05 / 0.8780 | -0.26 / 0.3918 | 0.2062 |
| Fucose w/o S | -0.11 / 0.6694 | -0.51 / 0.1252 | -0.26 / 0.3111 | -0.08 / 0.8020 | -0.26 / 0.3558 | 0.1932 |
| **Active TB** |  |  |  |  |  |  |
| Sialic Acid | 0.16 / 0.6853 | -0.21 / 0.6004 | 0.73 / 0.6406 | -0.48 / 0.7478 | -0.12 / 0.7744 | 0.9601 |
| Galactose | 0.01 / 0.9879 | -0.26 / 0.5207 | 0.53 / 0.7462 | -0.34 / 0.8304 | -0.13 / 0.7662 | 0.9745 |
| Fucose | -0.23 / 0.5701 | -0.09 / 0.8499 | 0.73 / 0.6292 | -0.60 / 0.6939 | -0.06 / 0.8827 | 0.9435 |
| GlcNAc | -0.22 / 0.5759 | -0.09 / 0.8613 | 0.34 / 0.8182 | -0.37 / 0.7994 | -0.05 / 0.9159 | 0.9446 |
| Fucose w/ S | -0.04 / 0.9107 | -0.24 / 0.5816 | 0.51 / 0.7306 | -0.35 / 0.8132 | -0.12 / 0.7886 | 0.9735 |
| Fucose w/o S | -0.18 / 0.6371 | -0.21 / 0.6033 | 0.74 / 0.6299 | -0.49 / 0.7423 | -0.12 / 0.7816 | 0.9543 |

Std. $\beta$ – Standardized Beta from multiple linear regression

